# Octopamine affects gustatory responsiveness and associative learning performance in bumble bees

**DOI:** 10.1101/2022.06.21.497037

**Authors:** Felicity Muth, Emily Breslow, Anne S. Leonard

## Abstract

Octopamine has broad roles within invertebrate nervous systems as a neurohormone, neurotransmitter and neuromodulator. It orchestrates foraging behavior in many insect taxa via effects on feeding, gustatory responsiveness and appetitive learning. Knowledge of how this biogenic amine regulates bee physiology and behavior is based largely on study of a single species, the honey bee, *Apis mellifera*. Until recently, its role in the foraging ecology and social organization of diverse bee taxa had been unexplored. Bumble bees (*Bombus* spp.) are a model for the study of foraging and learning, and its neural basis, but whether octopamine similarly affects sensory and cognitive performance in this genus is not known. To address this gap, we explored the effects of octopamine on sucrose response thresholds and associative learning in *Bombus impatiens* via conditioning of the Proboscis Extension Reflex (PER) using a visual (color) cue. We found that octopamine had similar effects on bumble bee behavior as honey bees, however, higher doses were required to induce these effects. At this higher dose, octopamine lowered bees’ sucrose response thresholds and appeared to enhance associative learning performance. Adding to recent studies on stingless bees (Meliponini), these findings support the idea that octopamine’s role in reward processing and learning is broadly conserved across Apidae, while pointing towards some differences across systems worth exploring further.

## Introduction

Octopamine (OA) is a biogenic amine involved in a diverse suite of physiological processes in insects (Roeder, 1994; Roeder, 1999). In honey bees (*Apis mellifera*) it may influence phenomena as diverse as circadian and cardiac rhythms (Bloch and Meshi, 2007; Papaefthimiou and Theophilidis, 2011), the stress response (Harris and Woodring, 1992) and motor performance (Fussnecker et al., 2006). However its clearest role is in the nervous system where it mediates sensory and cognitive processes associated with feeding (Giurfa, 2006; Rein et al., 2013). Alongside other biogenic amines (e.g. Dopamine (DA) and Tyramine (TA), OA’s precursor), OA has well-established effects on sensory responsiveness (Barron et al., 2002; Scheiner et al., 2014; Schilcher et al., 2021), including responsiveness to sucrose (Pankiw and Page, 2003; Scheiner et al., 2002). These effects on gustatory responsiveness are in turn a key determinant of learning performance in a foraging context (Scheiner et al., 2001). OA is centrally involved in the reward pathways that underlie appetitive learning: its injection into brain regions involved in learning and memory substitutes for a reward in a PER (Proboscis Extension Reflex) conditioning paradigm (Hammer and Menzel, 1998; Riemensperger et al., 2005; Schwaerzel et al., 2003; Unoki et al., 2005). OA’s heightened presence in the brains of starved foragers suggests that it also helps regulate the appetite—and perhaps more broadly, the motivation to learn—of workers in a feeding context (Mayack et al., 2019), (see also Akülkü et al., 2021).

These effects of OA on individual *A. mellifera* behavior may scale up to influence the division of labor and collective foraging efforts more generally (Wagener-Hulme et al., 1999). OA receptor expression in the brains of nurses vs. foragers differs (Reim and Scheiner, 2014; Schulz and Robinson, 2001), as do OA titers (Schulz et al., 2002). Among foragers, patterns of OA receptor expression change with age (Peng et al., 2021) and OA-mediated differences may underlie individual-level patterns of resource specialization (Arenas et al., 2021; Giray et al., 2007). For example, OA’s influence on sucrose response thresholds determines the quality of food they bring back when foraging (Giray et al., 2007; Pankiw and Page Jr., 1999). Pollen foragers have lower sucrose response thresholds and as such are less discriminating in the nectar they will accept compared to nectar foragers (Page Jr et al., 1998; Scheiner et al., 2001). OA also mediates social transmission of information about food resources: for example, bees treated with OA over-represent the quality of the forage they encounter when communicating with nestmates via their ‘dance language’(Barron et al., 2007). Interestingly, OA affects dances for both pollen and nectar quality in the same way, indicating that it plays a role in reward processing more broadly, and thus has an role equivalent to the dopaminergic system in mammals (Wise, 2004).

Given how clearly OA is involved in the regulation of individual and colony-level foraging behavior in *A. mellifera*, what role does it play for other bees? A 2022 Web of Science search of the scientific literature for “octopamine + bee” confirmed that while honey bees have historically offered a tractable model for untangling complex relationships between aminergic systems, individual physiology and collective behavior, other taxa are rarely considered (Fig. 1). Perhaps this reflects the assumption that OA’s involvement in these sensory and neural processes are so fundamental that they must be broadly conserved, though recent reviews highlight the need for more information across species (rev. Kamhi et al. 2017; Sasaki et al. 2021). Indeed, a recent study of the closely-related TA signaling system pointed towards a shared neural expression of TA receptors among representatives of Apini, Bombini, Meliponini, and Osmiini (Thamm et al., 2021), although behavioral data is needed to confirm if similar expression patterns relate to similar functionality. Likewise, behavioral work on stingless bees points to a conserved effect of OA on sucrose responsiveness and foraging behavior: *Melipona scutellaris* fed OA had a lower sucrose reponse threshold (Mc Cabe et al., 2017), and *Plebia droryana* foraged on a sucrose feeder containing OA at a faster rate compared to their behavior at a control feeder (Peng et al., 2020).

**Figure 1:**
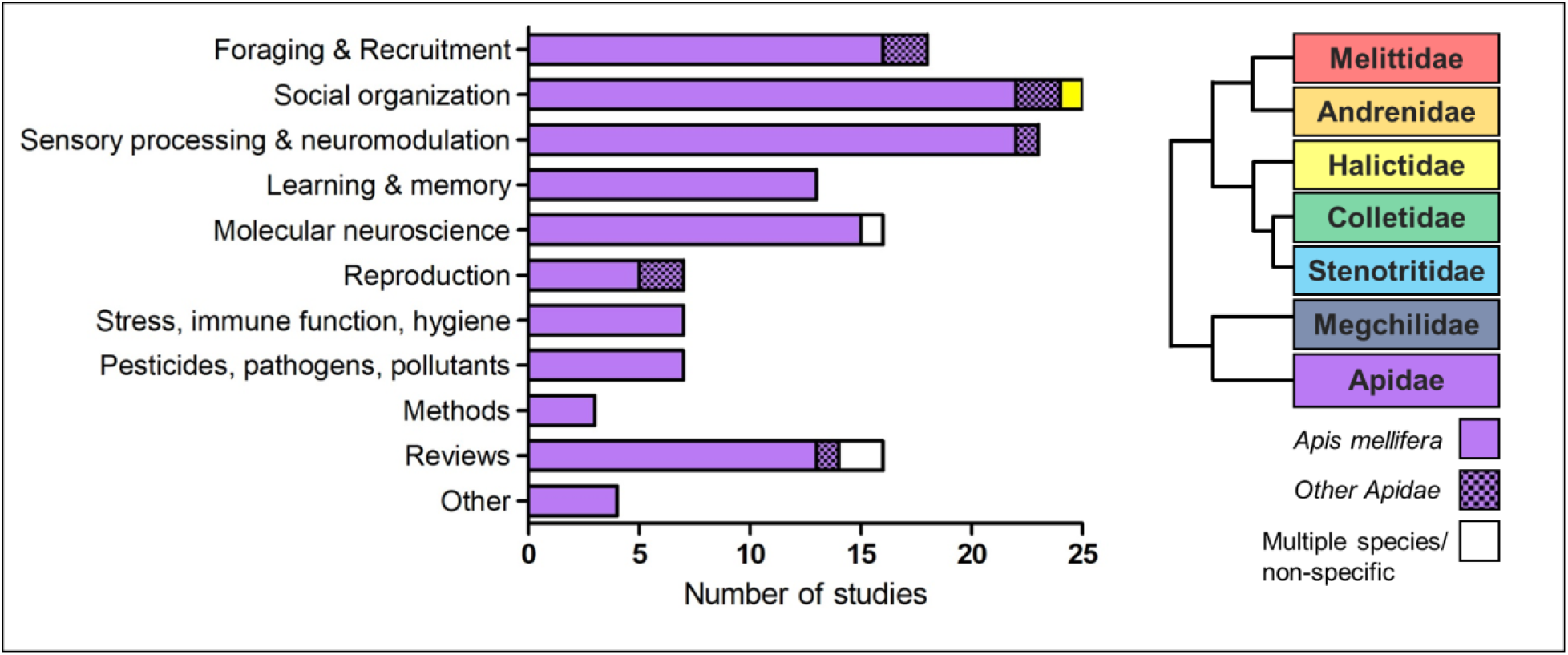
Summary of studies from a 2022 Web of Science search of the scientific literature for “octopamine + bee”. Color indicates bee family; Apidae and specifically *Apis mellifera* are greatly over-represented in the literature compared to other bee families.

On the other hand, recent comparative work has also revealed intriguing potential for differences in aminergic pathways. Thamm et al.’s (2021) study noted genus-level differences in the expression patterns of a tyramine receptor (AmTAR1) within the optic lobes. Likewise, within honey bees, OA receptor SNPs were associated with different ecotypes raising the prospect of their role in adaption to elevation-specific foraging ecologies (Wallberg et al., 2017). Given variation in bee sociality, dietary specialization and life histories (often involving both social and solitary foraging phases), exploring whether the behavioral effects of OA that are most established in *A. mellifera* manifest in other species will help fill in the picture of how this appetitive system supports diverse foraging behaviors across the bee tree of life.

Bumble bees (*Bombus*) are an important model for the study of insect cognition and foraging behavior (Chittka and Thomson, 2001). Like *Apis, Bombus* are generalist foragers that visit a variety of flowers when foraging, and as such must rapidly discriminate between floral rewards (e.g. nectars differing in sucrose concentration) and learn which flowers contain the highest quality rewards based on associated floral stimuli (color, scent etc.). Typically living as part of a colony, bumble bees communicate information about resource availability, albeit through chemical communication rather than a waggle dance (Dornhaus et al., 2003). Despite these shared features, bumble bees show a number of cognitive (Sherry and Strang, 2015), and neural (Gowda and Gronenberg, 2019) differences from honey bees. Given that individual *Bombus* workers are less specialized in their roles within the colony than in *Apis* and in their collection of resources more generally (Goulson, 2003), OA’s role in coordinating foraging-related behaviors is an open question.

Here we addressed the role of OA in bumble bee sensory responsiveness and cognition. Following a protocol similar to those used in the past with honey bees (Pankiw and Page, 2003; Scheiner et al., 2002) and stingless bees (*Melipona scutellaris*; Mc Cabe et al., 2017), we addressed how OA affected sucrose responsiveness and learning of a visual association in bumble bees *B. impatiens*. If OA has a similar role in bumble bees as it does in honey bees and stingless bees, then we expected its ingestion to lower sucrose response thresholds and enhance appetitive learning in a dose-dependent manner.

## Methods

### General methods

In all experiments we used *Bombus impatiens* workers (Experiment 1 n=65; Experiment 2 n = 56) purchased from Koppert Biological Systems (Howell, MI, U.S.A.). To obtain individuals for testing, we used an insect aspirator to remove bees from wicked feeders (Exp. 1: 30% (w/w) sucrose; Exp. 2: 15% (w/w) sucrose) in a central foraging arena (L × W × H: 100×95×90 cm) which had 3-5 colonies attached at any one time. We supplemented colonies with 5g of honey bee pollen (Koppert Biological Systems, Howell, MI, U.S.A.) every two to three days.

Following Riveros and Gronenberg (2009) and Riveros et al. (2020), we cooled bees in plastic vials placed on ice to immobilize them. Bees were then placed into individual plastic tubes (modified 1000 µl pipette tips, Fig. 2a) and restrained with two metal insect pins forming a “yoke” between their head and thorax that was secured with tape to the plastic tube (as in Muth et al., 2015; Riveros and Gronenberg, 2009). The bee could extend its proboscis and move its antennae but was otherwise immobilized. Bees were left to acclimate for three hours at room temperature in a dark room. After this time, we screened bees for responsiveness by presenting a droplet of 30% (w/w) sucrose to their antennae; bees that did not exhibit PER were removed from the experiment.

**Figure 2:**
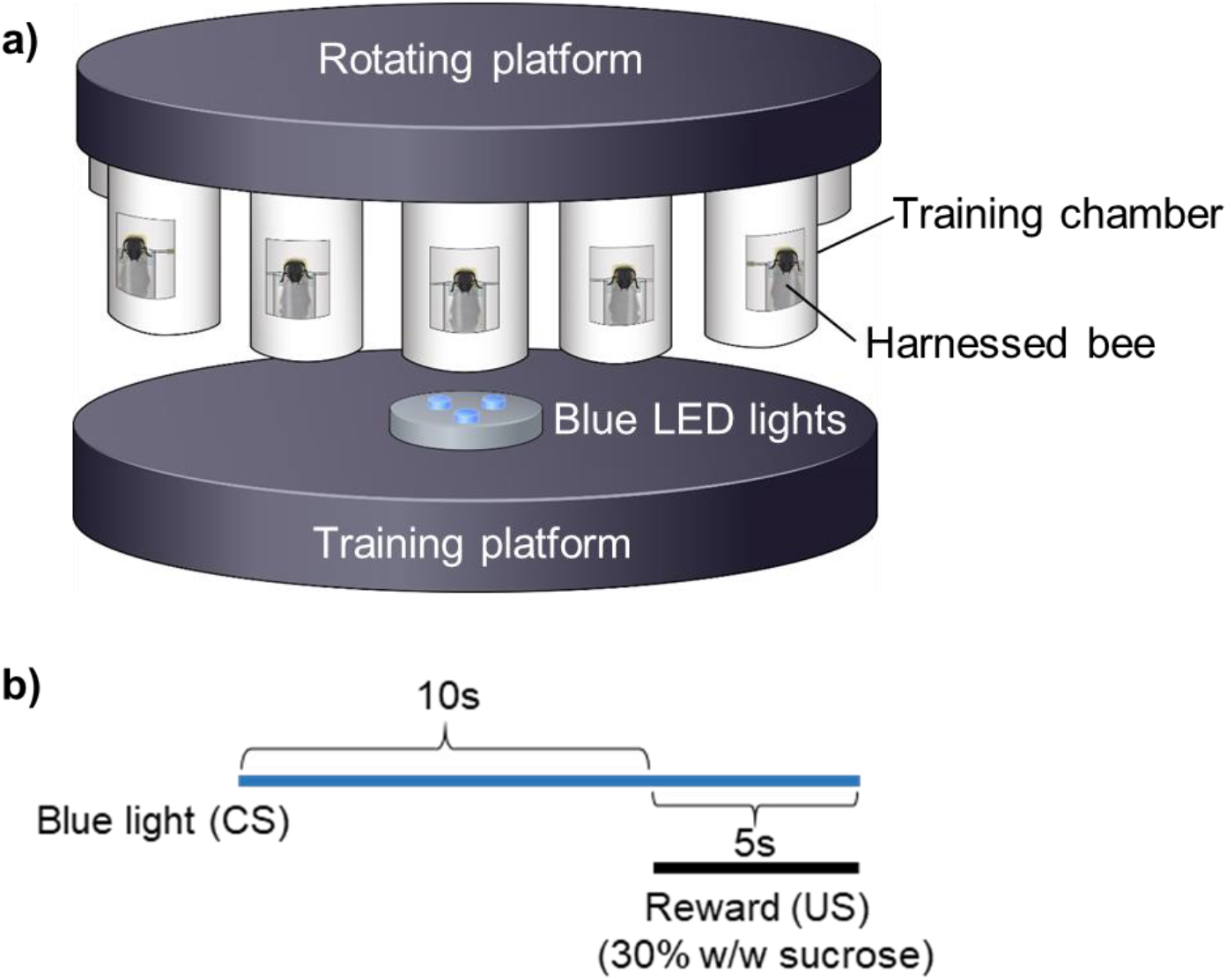
A diagram of the Proboscis Extension Response (PER) a) training apparatus and b) training protocol used in Experiment 2.

All experiments were conducted in a dark room, illuminated only with a red light to reduce any additional visual stimuli that could influence responsiveness or learning. Likewise, in all experiments, we fed bees OA, rather than injecting it. At least in honey bees, oral consumption has similar effects to injection but is less invasive (Barron, Schulz, & Robinson, 2002; Pankiw & Page, 2003)).

All statistical analyses were performed in R version 4.1.2 (2021) (R Core Team, 2020). We carried out GLMMs using the glmer function in the lme4 package; (Bates et al., 2015), including “bee” as a random factor to control for the multiple measures per bee. To determine the significance of interaction effects, we ran models with and without the interactions and used the anova() function to compare the fit of models using AICs. We carried out post-hoc tests using the emmeans package (Lenth 2017) and visualized relationships using effects() (Fox 2003).

#### Experiment 1: Does OA affect gustatory responsiveness in bumble bees?

To determine whether OA affected sucrose response thresholds, we assigned bees randomly to one of three treatments that varied in the solution they were fed prior to testing. In all treatments, we used a Hamilton syringe to feed bees 10µl of 30% (w/w) sucrose containing 1) 0µg/µl **OA** (control); 2) 2µg/µl **OA**; or 3) 8µg/µl **OA** (sample sizes in Fig. 3). After feeding bees, we allowed them to sit for 30 minutes to allow full absorption of the **OA** (Pankiw & Page, 2003). All three treatments were represented on a given day.

**Figure 3:**
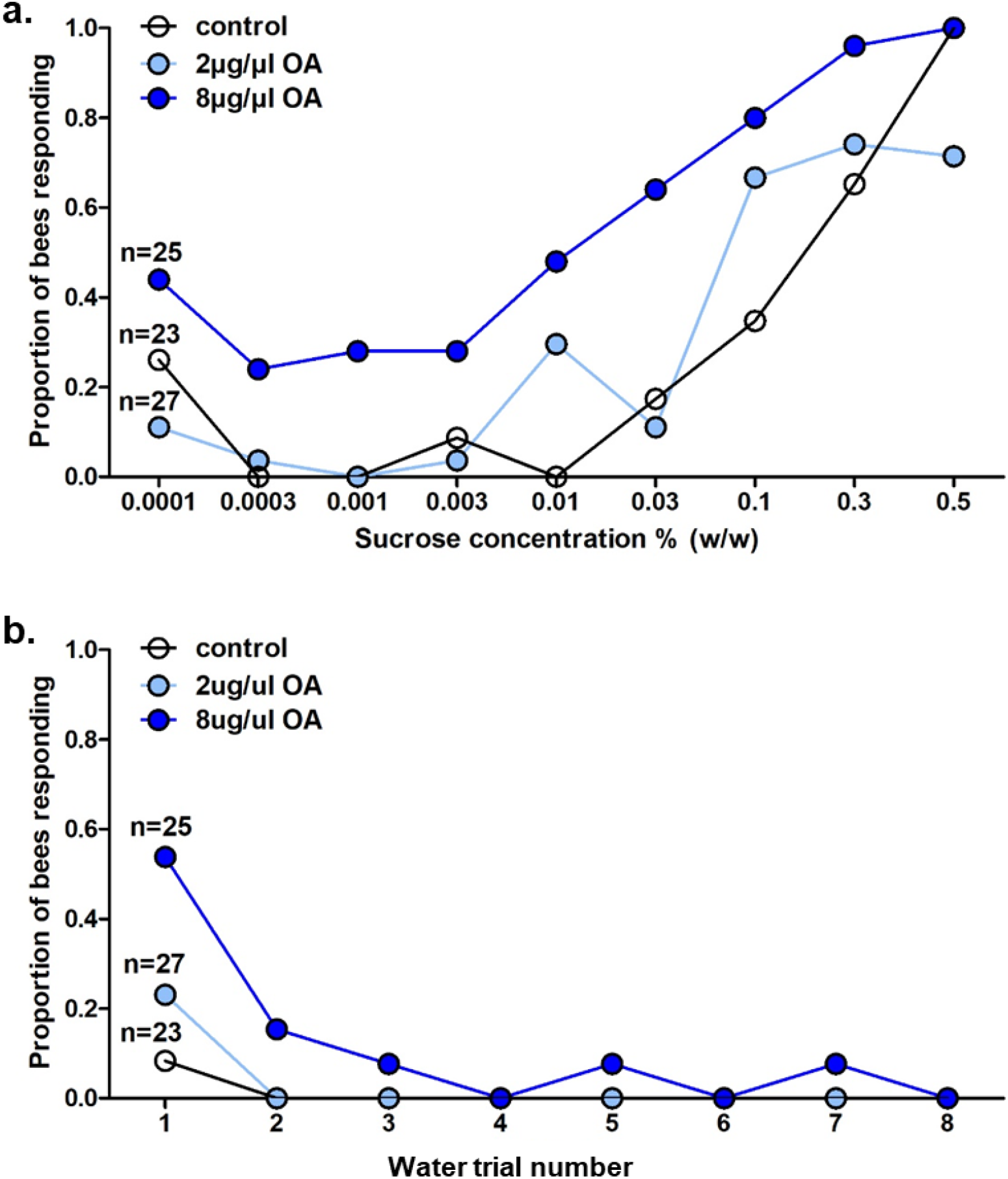
OA effects on bumble bee sucrose responsiveness (Experiment 1). When bees were pre-fed OA of two doses, a) sucrose responsiveness increased at the higher, but not lower, dose and b) initial responsiveness to water was higher in the high OA-treated group.

We tested the sucrose responsiveness of all bees by presenting them with eight different concentrations (w/w) of sucrose solution in succession (0.01%, 0.03%, 0.1%, 0.3%, 1%, 3%, 10%, 30%, 50%), with a presentation of water between each sucrose presentation (as in Mc Cabe et al., 2017; Pankiw and Page, 2003). As in these previous studies, presentation of water allowed us to distinguish a possible increase in sucrose responsiveness from a generalized increase in responsiveness to all stimuli. For each water trial, we presented the liquid to the bees’ antennae and allowed them three seconds to respond, before presenting them with the sucrose solution, and again giving them three seconds to respond. The inter-trial-interval between each sucrose presentation was 5 minutes.

#### Experiment 1 Data Analysis

To determine whether bees assigned to the three pre-treatments differed in their responsiveness to sucrose, we carried out a binomial GLMM with the binary response variable of whether the bee responded or not (1/0) and the following explanatory variables: sucrose concentration (continuous), treatment (3 levels) and the random factor “bee”. We initially planned to use a similar model to compare responsiveness to water, but due to the large number of bees not responding at all to this stimulus, we just compared the first water trial where there was the greatest response using a binomial linear model with the response variable responded or not (0/1).

#### Experiment 2: Does OA affect visual learning in bumble bees?

We harnessed 56 bees and trained and tested them using the proboscis extension response (PER) protocol. Bees were randomly assigned to two treatments, and fed prior to training 10μl of 30% (w/w) sucrose containing either 1) 0µg/µl **OA** (control; n=28) or 2) 8µg/µl **OA** (treatment; n=28). This dose was informed by our findings from Experiment 1. After being fed, individuals were transferred to the PER training apparatus and left to sit for 30 minutes before undergoing training and testing. Bees from both treatment groups were represented equally on each testing day.

The PER training apparatus consisted of a circular rotating platform suspended above the tabletop (Fig. 2a). Twelve ‘training chambers’ created from plastic cylinders were glued to the underside of this platform, approx. 6 cm apart. An opening (w×h: 3cm×1.5cm) in each training chamber allowed experimental access to the harnessed bee. Apart from a thin platform supporting the harnessed bee, the underside of each training chamber was open, allowing light to enter in from below (on which three blue (λ =470 nm) LED lights were mounted). Each chamber was lined with aluminum foil to evenly disperse lights which were controlled via a switchboard.

In an absolute conditioning paradigm, each bee was given 11 training trials followed by a test trial. Each training trial consisted of a presentation of the conditioned stimulus (blue light), followed by the unconditioned stimulus (30% (w/w) sucrose). In the initial trials, we exposed a bee to the light stimulus for 10 seconds before presenting the bee with the sucrose reward for an additional five seconds (2 seconds to antennae, 3 seconds to proboscis) (Fig. 2b). After the bee showed a conditioned response, the reward was presented (for 3 seconds) as soon as the bee extended its proboscis (even if 10 seconds had not elapsed). In all cases the reward and stimulus were removed simultaneously. As in Exp. 1, we used an inter-trial-interval of 5 minutes. The test trial was the same as the training trials with the exception that the blue light stimulus was given without the reward. In all learning and test trials we recorded (via live observation) whether the individual bee extended its proboscis in response to the blue light, and in cases when they did not but were presented with a reward (i.e. during the learning trials), if they responded to the presentation of the reward. This allowed us to not only determine if learning performance differed between the treatment groups but also if overall tendency to respond to sucrose presentation also differed.

#### Experiment 2 Data Analysis

If a bee did not exhibit a proboscis extension to presentation of the sucrose reward more than 4 times across the 11 training trials then we considered it to be unresponsive and excluded it from further analysis (OA n=1; control n=5), resulting in final sample sizes of OA n=27 and control n=23. To analyze whether bees learned differently across trials on the basis of treatment, we carried out binomial GLMMs where the response variable was whether the bee responded to the light stimulus or not (0/1) prior to receiving a reward, and the explanatory variables included were trial, treatment, and the random factor bee. Because both groups showed evidence of learning initially but then a decline in after trial 6, we split the data into two models: trials 1-6 and trials 7-11. The test trial data were analyzed alone using a binomial GLM.

To address whether feeding motivation/ responsiveness varied across trials we also carried out models, this time using all 56 bees tested. We included the response variable of whether the bee responded to the sucrose or not once it was presented to them (0/1) and the same explanatory variables as above. Interactions between trial and treatment were always included initially, but excluded if non-significant.

## Results

### Experiment 1: Does OA affect gustatory responsiveness?

Bees that were pre-fed the higher dose of **OA** were more responsive to sucrose than both the control and lower-dose treatment, which did not differ to each other (comparison of models with and without treatment × concentration interaction: χ^2^_2_ = 6.830; *p* = 0.033; Tukey post-hoc comparison between treatments: control vs. low: *z* = 0.761, *p* = 0.727; control vs. high: *z* = 4.713, *p* <0.0001; low vs. high: *z* = −4.302; *p* = 0.0001; Fig. 3a).

Similarly, in the first water trial, bees assigned to the high-dose pre-treatment were more responsive than the control group (*z* = 2.408, *p* = 0.016; Fig. 3b) while the bees that were pre-fed the lower dose of OA did not differ from the control bees (*z* = 0.103; 0.918; Fig. 3b). After the first water trial, bees across all treatments rarely responded at all.

### Experiment 2: Does OA affect visual learning in bumble bees? Learning performance – response to the conditioned stimulus

Across the first 6 learning trials, performance improved in both bees pre-treated with **OA** as well as in control bees (*z* = 4.731, *p* <0.0001) but the **OA**-treated bees showed higher performance (*z* = −2.196, *p* = 0.028). From the 7^th^ to 11^th^ learning trial, performance declined in both groups and there was an interactive effect, where the OA-treated bees at first out-performed the control group, but this effect disappeared towards the end of training (treatment × trial: *z* = 2.021; *p* = 0.043; trial *z* = −2.781; *p* = 0.005; treatment: *z* = −2.205, *p* = 0.027; Fig. 4a). There was no effect of treatment in the test phase (*z* = 0.167; *p* = 0.867), however overall response was very low by this point (Fig. 4a).

**Figure 4:**
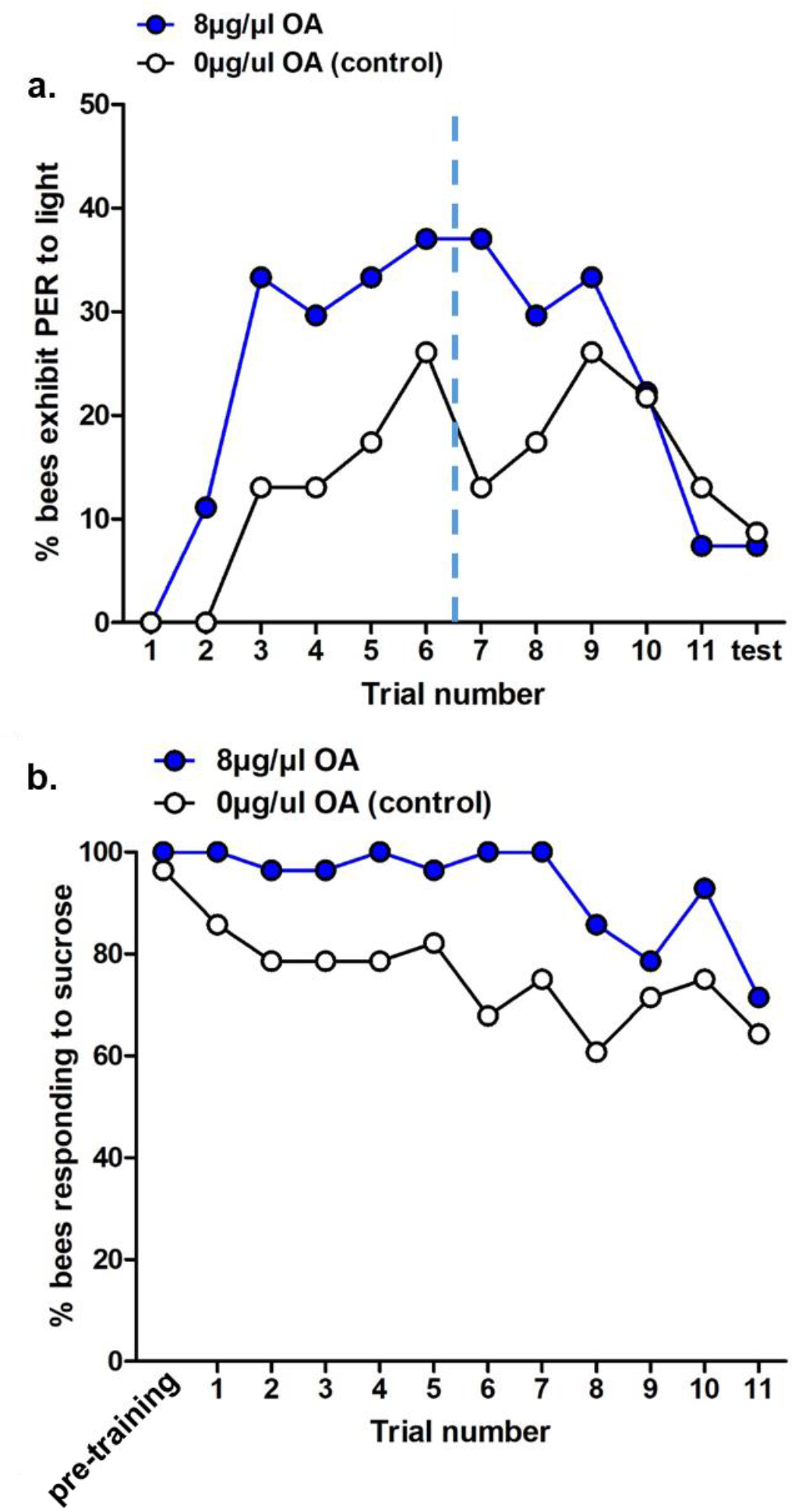
OA effects on bumble bee learning (Experiment 2). a) Bumble bees pre-fed a high dose of OA were more responsive to the conditioned stimulus than a control group; dashed line indicates where motivation to respond dropped across both treatments. b) The proportion of bees responding to the sucrose reward was higher in the OA-fed group than the control group.

### Responsiveness – response to the unconditioned stimulus

To address whether bees’ motivation to respond to the unconditioned stimulus (sucrose reward) varied across treatments, we compared whether bees in the OA-treated and control groups responded similarly once the sucrose reward was presented to them. Our results suggest that initially the motivation to feed dropped in the control treatment but remained in the OA treatment; however towards the end of the training period bees assigned to both treatments showed similarly low motivation to consume the sucrose reward (treatment × trial: *z* = 2.444; *p* = 0.015; trial *z* = −4.347; *p* < 0.001; treatment: *z* = −3.604, *p* < 0.001; Fig. 4b).

## Discussion

Octopamine (OA) has long been known to play an important role in honey bees (rev. Giurfa, 2006; Roeder, 1999), a system often used as a model to study the neural basis of behavior (Menzel, 2012) and the physiological mechanisms of task specialization (Riveros and Gronenberg, 2010). Yet, how OA affects behavior and physiology in other bee taxa exhibiting different levels of sociality (e.g. Halictidae: Jeanson et al., 2008; Smith et al., 2019); (*Ceratina*: Cook et al., 2019) is only beginning to be explored (Fig. 1). Our understanding of how OA mediates collective foraging in other social bees (e.g. Meliponinae; Mc Cabe et al., 2017; Peng et al., 2020) is equally limited. Within *Bombus*, only five prior studies have, to our knowledge, directly measured or manipulated OA. Four of these involve measuring OA levels or related gene expression with the aim of understanding reproductive division of labor: Bloch et al. (2000) found that OA titers in *Bombus terrestris* correlated with the dominance status of workers, independent of age or ovarian development; more recently Sasaki et al. measured OA levels in *Bombus ignitus* queens at different reproductive stages (Sasaki et al., 2017) or across workers vs. queens (Sasaki et al., 2021). Besides the present study, the only other experiment on *Bombus* that considers OA’s role in a foraging context appears to be Cnaani et al. (2003) which asked whether OA altered floral choice in *B. impatiens*. This experiment used a free-flying assay with automatically refilling artificial flowers to show that the presence of OA in “nectar” impacted *B. impatiens* workers’ persistence visiting a food source that became unrewarding.

Although these results have intriguing implications for understanding how nectar chemistry might activate octopaminergic pathways (Muth et al., 2022), this experiment was not designed to identify the mechanism behind shifts in floral choice. Indeed, understanding how OA (or other biogenic amines) influences foraging behavior in diverse bee taxa will require standardized and replicable behavioral assays. To this end, we adapted two protocols that have long been widely used to study the effects of OA on honey bee (and recently, stingless bee) learning. Using these, we found that OA has an analogous effect on bumble bees as in these two other genera, lowering sucrose response thresholds and enhancing associative learning. Our results indicate that similar mechanisms may underlie appetitive learning within Apidae, but also highlight differences that may inform future work in this and other systems.

Our first experiment explored how consumption of OA at two concentrations affected bees’ responsiveness to water and sucrose solutions. Broadly in keeping with work on honey bees, we report the first evidence that OA consumption increases sucrose responsiveness in *Bombus*. As in *Apis*, effects were dose-dependent: bees fed a higher dose of 10µl of 8µg/ µl (80µg total) were more responsive to sucrose across nearly all concentrations, and initially more responsive to water. Our lower-dose treatment (10µl of 2µg/ µl = 20µg total) were not more responsive to either stimulus type than the control bees pre-fed a control sucrose solution. Scheiner et al., (2002) assayed honey bees using a similar method and found analogous dose-dependency. In contrast to our findings with *Bombus*, honey bees in this previous work demonstrated a heightened sucrose responsiveness following exposure to much lower doses of OA (1.9 and 9µg). In a second study of OA’s effects on honey bees, increased sucrose responsiveness occurred following doses of 0.2, 2.0 and 20 µg (Pankiw and Page, 2003). In stingless bees, Mc Cabe et al. (2017) compared the sucrose responsiveness of bees following doses of 9.5, 19, and 38 µg OA and reported effects at the lowest doses as well. These differences in effectiveness of the lowest doses are unlikely to be due to differences in protocol, since in all these studies bees were immobilized and responsiveness was measured in a similar fashion. Without further data we cannot identify the source of this discrepancy. Body size is certainly a plausible explanation, but more subtle differences—for example, differences in receptor type or density—cannot be ruled out. As Mc Cabe et al (2017) noted, when OA is consumed by honey bees its behavioral effects are clear but their etiology is not: OA might change brain titers directly, or via more complex signaling cascades (as Scheiner et al, 2017 showed for TA). In addition to the dose difference noted here, discrepancies between *A. mellifera* and stingless bees in the timing of OA-enhanced sucrose responsiveness were noted by Mc Cabe et al (2017) raising the prospect that OA may exert its effects on sucrose responsiveness differently across taxa.

Also in keeping with previous findings from honey bees, we found that when we used the higher dose of OA (80µg) in Experiment 2, pre-consumption of OA enhanced learning performance, at least during the acquisition phase. While the PER protocol carries the advantage of being able to tightly control stimulus and reward presentation, it is limited in that the only behavior that is recorded is the bees’ tendency to extend its proboscis, which can be confounded with factors aside from learning and memory (rev. Muth et al., 2017). Several mechanisms could thus give rise to this effect. First, although we attempted to control for motivational effects by removing bees that did not respond to sucrose before starting the learning trials and by excluding bees that did not respond to sucrose more than 4 times across the 11 trials, there were still clear differences in motivation between the two groups (Fig. 4b). Namely, over the course of all trials, OA-fed bees were more likely to extend their proboscis to consume the sucrose reward than control bees (i.e. they showed a differential response to the unconditioned stimulus). As such, the differences seen between the treatments in bees’ tendency to extend their proboscis towards the conditioned stimulus may reflect motivational differences as much as differences in learning aptitude.

Work from honey bees also suggested that OA may have had the capacity to affect sensory responsiveness to features of both the unconditioned stimulus (US+) and conditioned stimulus (CS+) in ways that could promote learning performance. For example, given that Exp.1 established clear effects on sucrose responsiveness, bees in the treated group might have perceived the value of the US+ as higher value than control bees, a feature that can boost learning performance. It is also possible that OA’s ability to increase visual responsiveness (Scheiner et al. 2014) rendered the CS+ more salient to OA-dosed subjects in some way. Further work would be required to pinpoint the driver/s of the apparent performance difference we detected. Going forward, the effects of OA on learning and memory in bumble bees may be better addressed in protocols where bees are free-moving and where motivation vs. learning can be more easily differentiated (e.g. as in Muth and Leonard, 2019). While data collected similarly on this apparatus did not detect changes in responses through 8 training trials (Riveros et al., 2020) clearly our bees’ participation dropped markedly after the 6^th^ trial, due to satiation, fatigue, or other unknown factors. This led to few responses to the conditioned stimulus in the test phase across both groups, making them difficult to compare and likely obscuring any potential differences.

## Conclusion

Following OA consumption, results found in *Bombus* mirror those reported in *Apis* and *Meliponinae* in relation to sucrose responsiveness (both genera) and learning performance generally (which has only been measured in *Apis*). Yet, we did note some differences—namely, *Bombus* workers were not affected by our lower dose of OA, which work on the two other genera would have predicted to increase sucrose responsiveness. While subtle differences in OA-mediated behavior may not be significant for understanding broad patterns of aminergic-mediated social organization, we believe they are worth noting for two reasons. First, small changes in appetitive signaling pathways could be meaningful for understanding mechanisms involved in ecological radiation (Ji et al., 2020; Pankiw, 2003) as OA is clearly involved in determining what bees choose to collect and their motivation to do so. Secondly, many popular pesticides target OA receptors (Ahmed and Vogel, 2020; Farooqui, 2013; Papaefthimiou et al., 2013) and the OA signaling pathway in particular has been implicated in mediating bees’ responses to stress (Chen et al., 2008; Corby-Harris et al., 2020), pathogens and parasites (Mayack et al., 2015; Spivak et al., 2003), and pollutants (Søvik et al., 2015). In an era of wild bee declines, understanding whether *A. mellifera* is indeed a representative model for anthropogenic influence on aminergic pathways more broadly is a pressing challenge.

## Statements and Declarations

### Funding

This work was supported by the National Science Foundation IOS-1755096 (ASL) and IOS-2028613 (FM).

### Competing Interests

The authors have no relevant financial or non-financial interests to disclose.

### Author Contributions

ASL conceived of the experiments; experimental design was planned with input from EB and FM. EB collected the data. FM analyzed the data and co-wrote the manuscript with ASL. All authors read and approved the final manuscript.

### Ethics Approval

While no ethical approval was needed we aimed to minimize potential suffering to bees through cold-immobilizing them prior to placing them in harnesses for the experimental protocol. Bees were euthanized via freezing.

## Notes

### Competing Interest Statement

The authors have declared no competing interest.

